# Levodopa impairs lysosomal function in sensory neurons in vitro

**DOI:** 10.1101/2024.09.29.614220

**Authors:** Oyedele J. Olaoye, Asya Esin Aksoy, Santeri V. Hyytiäinen, Aia A. Narits, Miriam A. Hickey

## Abstract

Parkinson’s disease (PD) is the second-most common neurodegenerative disease world-wide. Patients are diagnosed based upon movement disorders, including bradykinesia, tremor and stiffness of movement. However, non-motor signs, including constipation, rapid eye movement sleep behavior disorder, smell deficits and pain are well recognized. Peripheral neuropathy is also increasingly recognized, as the vast majority of patients show reduced intraepidermal nerve fibers, and sensory nerve conduction and sensory function is also impaired. Many case studies in the literature show that high-dose levodopa, the primary drug used in the treatment of PD, may exacerbate neuropathy, thought to involve levodopa’s metabolism to homocysteine. Here, we treated primary cultures of dorsal root ganglia and a sensory neuronal cell line with levodopa to examine effects on cell morphology, mitochondrial content and physiology, and lysosomal function. High-dose levodopa reduced mitochondrial membrane potential. At concentrations observed in the patient, levodopa enhanced immunoreactivity to beta III tubulin. Critically, levodopa reduced lysosomal content and also reduced the proportion of lysosomes that were acidic while homocysteine tended to have the opposite effect. Levodopa is a critically important drug for the treatment of PD. However, our data suggests that at concentrations observed in the patient, it has deleterious effects on sensory neurons that are not related to homocysteine.

**Simple Summary:** Parkinson’s disease (PD) is one of the most common chronic, degenerative brain diseases worldwide. Patients are diagnosed on the basis of slowness of movement and/or tremor and/or stiffness. However, many symptoms that are not movement related are now well recognized. Patients show changes in skin sensation, and the vast majority of patients show loss of sensory neurites, which enable sensation in skin. These changes in skin sensation occur prior to diagnosis; however, sensory issues may also be exacerbated by levodopa, an important drug used in the treatment of PD. Undoubtedly, levodopa is critical for the treatment of PD, but at high doses, it has repeatedly been shown to impair sensation in PD patients. Here, we show for the first time that high-dose levodopa impairs function of sensory neurons. Importantly, we also show for the first time that lysosomes, a critical organelle involved in recycling, are impaired by levodopa concentrations observed in patients. These data are important given the well-known lysosomal dysfunction observed in PD. Our data sheds light on how levodopa, the most important drug in the treatment of PD, may contribute to sensory deficits in PD.

## 1. Introduction

Parkinson’s disease (PD) is the most prevalent neurodegenerative motor disorder worldwide, and indeed, it is currently placed 11^th^ in the global burden of disorders affecting the nervous system, with DALYS (disability-adjusted life years) increased by 10% since 1990 [1]. It was only in 1960 that Hornykiewicz first published the loss of dopamine in striatum of patients with PD [2]. A prior study in the 1930s had shown that dopa decarboxylase converted levodopa to dopamine [3], and Hornykiewicz and colleagues very quickly moved forward to a clinical trial of levodopa to show beneficial effects of levodopa that were not seen with related molecules[4,5]. Levodopa, as a chronic oral drug for PD, was established by 1967 [6], fast becoming the mainstay of treatment and the treatment of choice among patients [7] and neurologists [8–10]. Undoubtedly, levodopa treatment of PD is a success story.

The diagnostic movement disorder of PD is largely due to the loss of dopamine neurons of the substantia nigra pars compacta (SNpc), a tiny nucleus in midbrain with a critical projection to striatum. Although the precise mechanism(s) underlying loss of motor control in PD remains unclear, this loss of the neurons removes a modulating input to striatal output neurons. Loss of dopaminergic neurons of the SNpc in animals also leads to movement disorders[11], which are well treated with levodopa [12,13], and some of the movement disorders of PD are reproduced pharmacologically in schizophrenic patients treated with dopamine antagonists[14]. The specific cause of cell death in PD also remains unclear, although risk factors for PD include traumatic brain injury[15] and pesticides such as rotenone and paraquat [16], which are complex I inhibitors that lead to oxidative stress and mitochondrial impairment [17]. Genetic causes of PD are rare but very informative and include mutations and multiplications of the *SNCA* gene, which encodes alpha synuclein. In aggregated form, alpha synuclein is the majority protein in the Lewy body, a hallmark of PD neuropathology[18]. This protein is normally located in the vertebrate presynaptic nerve terminal, with a weak association to synaptic vesicles and a modulatory role in the release of dopamine[19], although many questions remain over its function also.

The clinical use of levodopa in the treatment of PD motor dysfunction has changed little – patients take several doses over the course of the day, due to the drug’s short halflife[10]. As the disease progresses, the frequency of dosing increases, but eventually, possibly due to the continual loss of dopaminergic synapses in striatum, patients experience the phenomena of “wearing-off”, whereby motor symptoms appear at the end of each dosing window – this is a significant problem for patients [9]. In addition, dyskinesias (involuntary, writhing-like movements) are well known and are linked to the combination of PD with the long-term, pulsatile use of levodopa[9]. Data suggesting a more continuous plasma level is beneficial in reducing these motor complications have existed for many years [20], and a very recent, large Phase III trial proved this point [21].

Nevertheless, PD is not just a movement disorder. It has long been known that peripheral signs, such as constipation, and olfactory impairment are observed in patients many years prior to the diagnostic movement dysfunction[8]. Moreover, skin, the largest organ of the body, is now becoming a major player in PD. In a very large, recent population study, skin symptoms were found to rise in the years prior to diagnosis of PD [15]. The vast majority of patients show loss of intraepidermal nerve fibres ([22]; IENFs) – the small fibres in skin epidermis, and the loss of IENF progresses with disease[23,24]. The cell bodies of skin sensory (nociceptive) neurons reside in dorsal root ganglia, which carry Lewy bodies from early stages in PD[18]. Symptoms of polyneuropathy are highly prevalent in patients [25], and are seen at diagnosis [26] and prior to diagnosis of PD [27].

Importantly, sensory deficits impact motor function by affecting the patient’s ability to gauge balance [28] and indeed, the Unified PD rating scale correlates with IENF loss[29]. Certainly, issues with gait and balance are clearly identified as refractory to current treatments and are a matter for urgent therapeutic development [10]. Thus, patients show several ongoing changes in skin, changes in skin sensation appear prior to the motor signs that are used for diagnosis, and disease-induced impairment of sensory neurons may underlie important symptoms that are currently undertreated in the clinic. Moreover, skin measures may be able to quantitatively and objectively monitor disease, particularly as skin-punch biopsies are safe and have a very low prevalence of side effects [30].

However, high-dose levodopa itself may also contribute to the neuropathy that is observed in PD. In the clinic, high-dose levodopa has repeatedly been associated with exacerbation or development of neuropathy symptoms [10,22] and has even been shown to impair spinal conduction in as little as one month [31]. Moreover, loss of epidermal sensory neurites (IENF) correlates with oral levodopa dose [29].

Here, in this paper, we show for the first time that levodopa is deleterious to primary cultures of dorsal root ganglia and a sensory neuronal cell line. We show that levodopa, at concentrations observed in patients, exacerbates toxicity induced by the pesticide rotenone, suggesting that in vivo, levodopa may add to parkinsonian pathophysiology. Thus, our highly translationally relevant data show that levodopa is deleterious to sensory neurons, and that treatment with entacapone does not prevent these deleterious effects on lysosomes. Our data suggest that levodopa may contribute to peripheral neuropathy in PD patients.

## 2. Materials and Methods

### 2.1 DRGs

#### 2.1.1 DRG preparation

Sprague Dawley pups (P5-P14) were euthanised by decapitation and dorsal root ganglia (DRGs) isolated and placed in ice-cold sterile PBS. DRGs were transferred to a dissociation solution (2 mg/ml collagenase, + 0.1 mg/ml DNase; Gibco) and incubated for 40 min at 37°C. Cells were then incubated in trypsin (0.05%) for an additional 5 min at 37°C and then in DMEM/F-12 + 10% foetal bovine serum (SigmaAldrich) to quench trypsinization. Cells were then triturated, centrifuged (10 min at 600 *g*) and the resulting pellet was resuspended in fresh complete medium (Neurobasal-A (Gibco) supplemented with B-27 (Thermofisher), gentamicin and glutamine (GlutaMAX, Thermofisher)). An equivalent of 2 ganglia per 50μl were plated per quadrant of 4-quadrant dishes (Cell Vis 4-chamber glass-bottom dishes; Gerasdorf Austria). Coverslips were coated with laminin and poly-D-lysine. After 3-4 hours at 37°C and 5% CO_2_, cells were supplemented with complete medium (Neurobasal A (ThermoFisher), B27 supplement (ThermoFisher), Gentamicin and glutaMAX (Thermofisher)). The following day, medium was replaced with fresh medium containing 1.5μM cytarabine (Ara-c; SigmaAldrich) to reduce proliferation of fibroblasts.

#### 2.1.2 DRG treatments

At approximately DIV 7, cells were placed into a hypoxia chamber (hypoxia: 3% O_2_, 5% CO_2_ and 92% N_2_) for 3 days to habituate. After 3 days, the cells were treated with rotenone (SigmaAldrich, 0, 1nM, 10nM, 100nM, 500nM) and/or levodopa methyl ester (0, 3μM, 30μM, 300μM) for 24 hours or 7 days. Hypoxia was used to prevent auto-oxidation of levodopa[32]. Vehicle controls for rotenone and levodopa were DMSO (Sigma Aldrich) and distilled water, respectively. For some oxidative stress experiments, additional cells were incubated in normal atmosphere (normoxia, 5% CO_2_, 20% O_2_) and were treated in parallel.

#### 2.1.3 Measurement of mitochondrial membrane potential

Membrane potential was measured using TMRM (Thermofisher) in hypoxic conditions only. Medium was removed and replaced with medium containing 10 nM TMRM (non-quenching[33,34]) and treatments (see above) for 30 min. Cells were then imaged using a confocal microscope (LSM780, ex 561nm, em 566-669nm; 0.05 × 0.05μm per pixel, 53.14 × 53.14μm total field of view taken using Plan-Apochromat 40×/1.3 oil DIC M27 objective). Lasers settings were consistent within each experiment. The photomicrographs were analysed using ImageJ software. Briefly, each soma was outlined to create an ROI and the mean pixel intensity of TMRM-stained mitochondria measured. 24hrs: N=3 experiments, N=9-19 cells per condition per experiment, 32-41 cells per condition overall. 7 days: N=6 experiments, N=4-60 cells per condition per experiment, N=99-173 cells per condition overall.

#### 2.1.4 Oxidative stress assay

Dihydroethidium (DHE; Santa Cruz Biotechnology) was used to measure reactive oxygen species (ROS) in live cells. Briefly, DHE was added to cells (final concentration 10μM DHE) and were incubated for 30 minutes. Cells were then imaged using a confocal microscope (LSM780, ex 561nm, em 585-733nm; 0.83 × 0.83μm per pixel, 1024 × 1024 pixels, taken using Plan-Apochromat 10×/0.45 M27 objective). Subsequently, the images were analysed with ImageJ using particle analysis. Briefly, images were thresholded using manual intensity threshold boundaries of 120-255. Particle sizes 200μm^2^-infinity in size and with a circularity of 0.10-1.00 were measured, which ensured that only DRG neurons were quantified and avoided the fibroblasts present in our mixed cultures (cultures were treated with Ara-c but were considered mixed as some fibroblasts remained) providing an outcome of percent area above threshold per image. Normoxia 24hrs treatment: N=1-8 images per condition per experiment, N=4 experiments and N=16-24 images per condition in total. Hypoxia 24hrs treatment: N=1-15 images per condition per experiment, N=4 experiments and 13-30 images per condition in total. Hypoxia 7 days treatment: N=3-40 images per condition per experiment, N=5 experiments and 74-142 images per condition in total.

#### 2.1.5 Immunostaining

Cells were fixed for 10 minutes using 4% paraformaldehyde (Sigma Aldrich) containing 250mM sucrose (Fisher Bioreagent) and stored at -4° C for subsequent immunostaining. Following washing (3 × 5mins in 0.01M PBS), cells were then permeabilised with 0.1% Triton X-100 for 10 minutes and then blocked in 5% goat serum (diluted in 0.01M PBS: block, Jackson ImmunoResearch) for 1 hr. Cells were then incubated in primary antibodies, diluted in block, overnight with gentle rotation (anti-MAP2, Abcam cat# ab5392 RRID:AB_2138153, 1 : 10 000; anti-beta III tubulin antibody, Abcam Cat# ab18207, RRID:AB_444319, 1 : 2500; anti-ATP5b, Millipore Cat# MAB3494, RRID:AB_177597, 1 : 500). Cells were washed (3 × 5mins in 0.01M PBS) and then incubated with secondary antibody for 2 hr (Jackson ImmunoResearch, 1:200 in block). Following washing (3 × 5mins in 0.01M PBS), cells were counterstained with 0.5 μg/ml Hoechst solution (Hoechst-34580 dye, Sigma Aldrich) for 10 minutes, then washed in PBS. Finally, a drop of fluorescence mounting medium was added to each quadrant, and dishes were then stored at -20°C until imaging.

Neurites based upon MAP2 staining were imaged using LSM780 AxioObserver, plan-apochromat 10×/0.45 M27 objective, image resolution 1.38×1.38μm/pixel, 1024×1024 pixels/image. Excitations and emissions were 405nm, 410-585nm (Hoechst), 488nm, 490-594nm (MAP2). 4-13 images were taken per treatment per timepoint per experiment. Percent area positive for MAP2 staining was determined following auto-thresholding in ImageJ. Data were then expressed as percent of control-treated cells at each timepoint. There were 1-14 photomicrographs per condition per experiment, N=4 experiments and 10-51 images per condition in total.

Neurites based upon beta III tubulin staining were imaged using LSM780 AxioObserver, plan-apochromat 10×/0.45 M27 objective, image resolution 1.66×1.66μm/pixel, 512×512 pixels/image. Excitations and emissions were 405nm, 410-499nm (Hoechst), 561nm, 585-733nm (beta III tubulin). Channels were separated, the red channel was autothresholded and then percent area above threshold per image was quantified in ImageJ (Fiji, 1.54f [35]). Using the thresholded images to “IdentifyPrimaryObjects”, fluorescence intensity per neurite per image was quantified from the red channel (beta III tubulin) in CellProfiler (V2.4.6 [36]). 20-40 images per condition from N=3 experiments.

Cell bodies were imaged using LSM780 AxioObserver, plan-apochromat 40/1.3 oil DIC M27 objective, image resolution 0.05×0.05μm/pixel, 1024×1024 pixels/image. Excitations and emissions were 405nm, 415-572nm (Hoechst), 488nm, 490-594nm (ATP5b), 561nm, 585-733nm (beta III tubulin). The images were then analyzed in ImageJ[35]. To determine mitochondrial content in soma, channels were split and each soma was outlined from beta III tubulin staining, avoiding the nucleus. Staining for ATP5b was autothresholded, and the percent area above threshold, within the soma ROIs, was quantified and expressed relative to control-treated cells. Control cells that were incubated with no primary antibody showed no staining. There were 5-66 cells per condition per experiment, N=3 experiments and 130-131 cells per condition in total.

#### 2.1.6 Lysosome content and acidity measurements

Following treatment with control or 300μM levodopa in hypoxia, medium was removed from cells and replaced with medium containing treatments and Lysotracker red (100nM, L7528 Thermofisher) and CellTracker green (6μM, C7025, Thermofisher). Cells were incubated in the dark, at 37°C, for 1 hour, washed gently and imaged in warmed Krebs. A 10-μm stack through the middle of each DRG soma, identified based upon CellTracker, was taken using an LSM780 confocal Plan-Apochromat 63×/1.4 Oil DIC M27 (pixel size 0.7μm × 0.7μm × 0.34μm). Using a beam splitter, a line scan was taken, ex405nm em410-556nm (Lysosensor), ex561nm em 566-691nm (LysoTracker). For analysis, channels were split, auto-thresholded and despeckled. A 5μm block of soma was outlined and surrounding area cleared. For total lysosomal content, area stained by Lysotracker per optical slice was quantified, expressed as a proportion of cell area and a mean per soma generated. For acidity, the proportion of Lysosensor contained with Lysotracker-stained ROIs per slice was quantified and a mean per soma generated. 28-40 soma were analysed per condition per experiment, N=3 experiments with 98-104 cells per condition in total.

### 2.2 50B11 cells

#### 2.2.1 Culturing

50B11 cells were cultured in Neurobasal medium supplemented with 10% fetal bovine serum, 2% B-27, 20mM D-glucose and 0.2 M L-glutamine (GlutaMAX). All media and supplements were from Thermo Fisher Scientific, USA. Cells were used at passages 5-9.

#### 2.2.2 50B11 treatments

Cells were plated in poly-D-lysine-coated 4-quadrant dishes (Cell Vis 4-chamber glass-bottom dishes; Gerasdorf Austria). The following day, they were placed in a hypoxic chamber (3% O_2_, 5% CO_2_ and 92% N_2_) for 24hrs to habituate. Cells were then treated with combinations of levodopa methyl ester (a more water-soluble version of levodopa, SigmaAldrich; 30μM, 300μM), entacapone (1μM [37]), homocysteine (20μM [38,39]) or vehicle (ddH_2_O) for 24hrs.

#### 2.2.3 Lysosome analysis

Following 24hrs of treatment, medium was replaced with medium containing treatments and Lysotracker red (100nM, L7528 Thermofisher Scientific) and Hoechst-34580 dye (500nM; Sigma Aldrich) for 30 minutes. Cells were then imaged using StereoInvestigator (V5.00, MBF Bioscience) on a Zeiss Z1 microscope at ×63 magnification. Lysotracker red is a well-known dye that is ion-trapped in acidic environments. Images of nuclei and lysosomes were taken separately through the depth of each cell using consistent settings for exposure, gain and binning within each experiment. The image stacks were then rendered using Helicon Focus (V8.2.2, Kharkiv, Ukraine) and cropped to 1550 × 1180 pixels. Resultant images were then processed in CellProfiler (V4.2.6; [36]). The size of all lysosomes was quantified from 7-24 images per condition per experiment, N=4 experiments and a final number of 43-77 images per condition in total (20275-37932 lysosomes in total per condition). In addition, Hoechst-stained nuclei were co-localised within these photomicrographs in CellProfiler and then expanded to create masks using boundaries of 10, 20, 30, 40, 50 and 60 pixels from original size. Lysosomes within each boundary were identified. To create total content per boundary per cell, the total area occupied by lysosomes per boundary per image was divided by the total area of nuclei per image. As above, 7-24 images per condition per experiment, N=4 experiments and a final number of 43-72 images per condition (5 lysosome photomicrographs for the homocysteine condition were not used as nuclei photomicrographs were unavailable).

### 2.3 Statistics

All analyses were conducted blinded and were then unblinded for statistical comparisons and graphing, which was performed using GraphPad Prism V10.2.3. The threshold for significance was p values < 0.05. Where variances were similar (based upon F tests), Student ttests were used to compare two independent groups and 1-way ANOVAs were used to compare between groups varying by 1 factor. Where data showed significantly different variances, based upon F tests, unpaired ttests with Welch’s correction were used to compare two independent groups, and Brown-Forsythe ANOVA tests were used to compare between several groups varying by 1 factor, followed by Dunnett’s T3 multiple comparisons test. Two-way ANOVAs (mixed-effects model) followed by appropriate post-hoc tests were used to compare data where there were two factors – equal variances were not assumed and the Geisser-Greenhouse’s epsilon correction was used. For large datasets (lysosome sizes), Matlab[40] was used for 2-way ANOVA using the p=ANOVA2 function. All conditions must have the same group size to use this function: a random non-repeating selection of 20000 lysosome sizes were taken from each condition using an Excel macro. Datasets on individual lysosome sizes were tested for normality using D’Agostino & Pearson and Kolmogorov-Smirnov tests. For Kruskal Wallis 1-way ANOVA analysis of this dataset, the full dataset was used. For graphs, data are shown as scatter plots showing individual technical replicates with mean ± sem or as box and whisker plots of technical replicates with 95% confidence intervals, with remaining datapoints, medians and means shown, as recommended [41–43]. In all cases, experimental means are superimposed for reference.

## 3. Results

### 3.1 Establishing a parkinsonian primary sensory neuronal cell model

We began our experiments with a preliminary study on the concentration of rotenone for inducing parkinsonism in primary cultures of dorsal root ganglia. Cultures were incubated in normoxia (21% O_2_) for these experiments. There was a consistent significant decline in MAP2 immunostaining by 4 days, at 1μM rotenone, and by 7 day at 500nM, primarily of neuronal processes (Figure 1; effect of treatment F (5, 178) = 21.3, p<0.0001).

**Fig 1.**
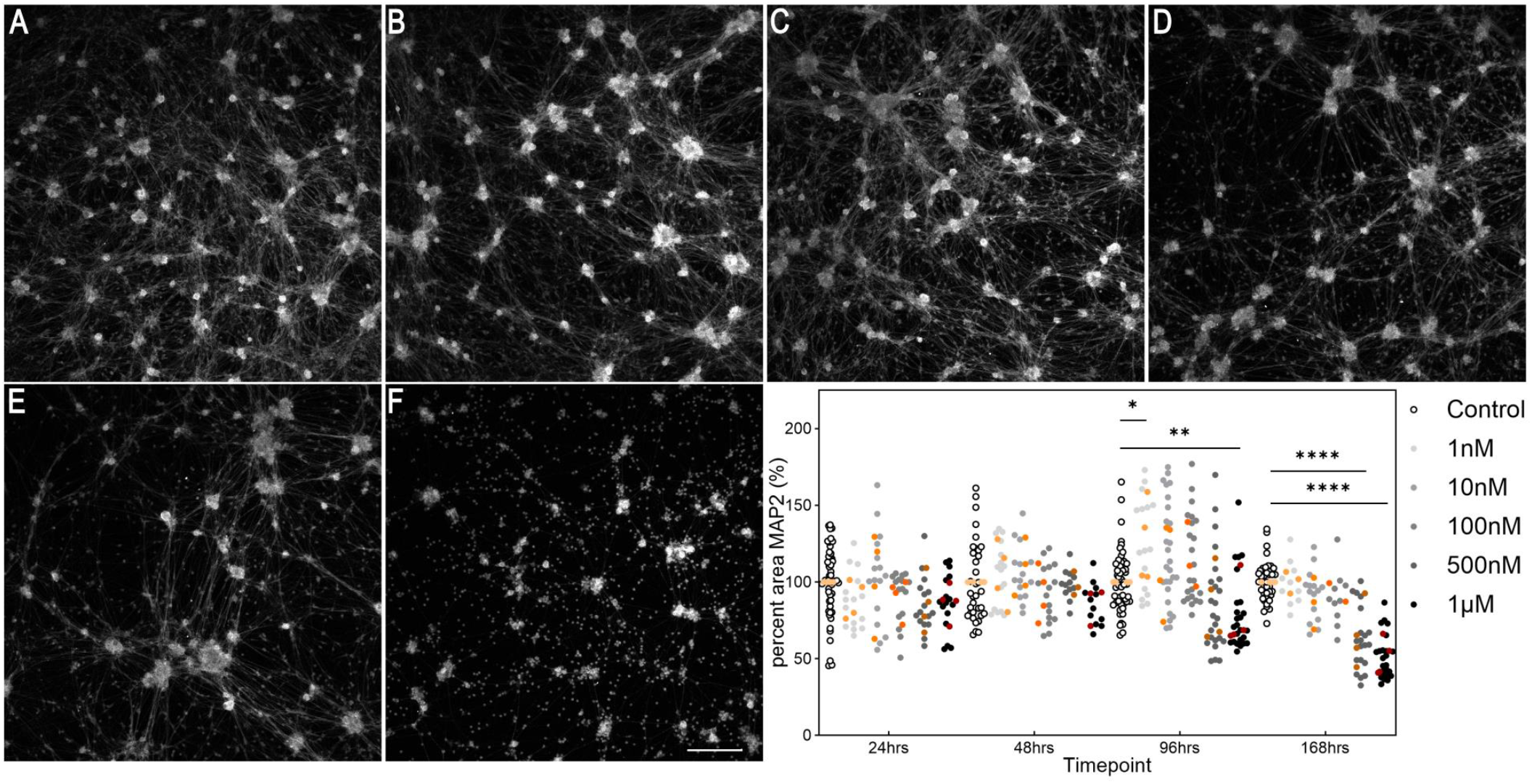
Time-dependent cytotoxicity of rotenone in DRGs. Cells were fixed and stained for MAP2, a pan-neuronal marker. Example photomicrographs following 7 days (168hrs) of treatment; A = control, B = 1nM, C = 10nM, D = 100nM, E = 500nM, F = 1μM. Scalebar in F = 200μm and is for all photomicrographs. Photomicrographs are not edited except for contrast, and they depict the MAP2 channel. The graph shows the time-dependent effect of rotenone on DRGs. Symbols in white and shades of gray to black show data from individual technical replicates. Symbols in shades of orange to brown show experimental means. *p<0.05, **p<0.01, ****p<0.0001

### 3.2 Effect of levodopa on sensory neurons alone and in the context of parkinsonism

#### 3.2.1 Levodopa exacerbates mitochondrial impairment in parkinsonism

Levodopa oxidises upon exposure to air[32] and in any case, the partial pressure of oxygen within the body is typically less than 5%[44]. Thus, culturing in a normoxic environment is actually hyperoxic with respect to endogenous conditions. Therefore, in order to examine the legitimate effects of levodopa on sensory neurons, we cultured in hypoxic conditions (3% O_2_). Although this is normoxic compared to within the body, for simplicity, we refer to 3% O_2_ as hypoxia/hypoxic. In vivo, in human patients, levodopa plasma levels fluctuate with each dose but concentrations vary from 1 – 10 mg/L [45–50]. These concentrations correlate to approximately 2-30μM, and here, we treated with 3, 30 and 300μM.

Levodopa has previously been shown to impair mitochondrial function in mesencephalic cultures and cancer cell lines[32] in hypoxia. However, here, in primary sensory neurons, we found that after 24hrs treatment, 30μM levodopa increased ΔΨ_M_, suggesting hyperpolarization in hypoxic conditions (Figure 2 A). By 7 days, this effect was lost and instead 300μM levodopa reduced mitochondrial membrane potential by 32% (Figure 2A; effect of levodopa, F (2.91, 530.1) = 10.7, p<0.0001, levodopa × time interaction F (3, 547) = 8.4, p<0.0001).

**Fig 2.**
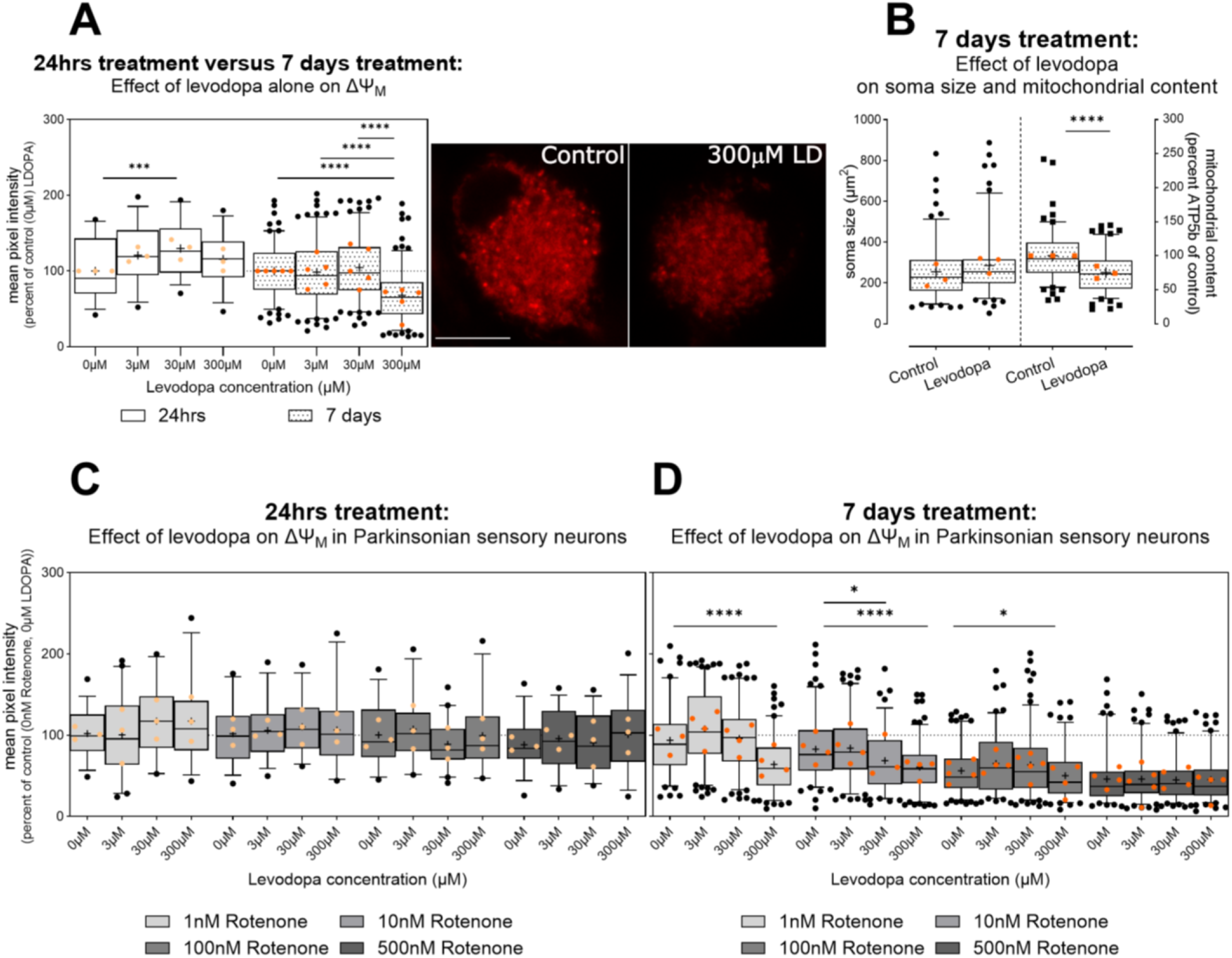
Effect of levodopa on mitochondrial membrane potential (ΔΨ_M_) in primary sensory neurons (DRGs) cultured in hypoxia. Data are normalized to percent of control cells treated with 0μM levodopa and 0nM rotenone in normoxia or in hypoxia. A) 30μM levodopa increases ΔΨ_M_ at 24hrs in hypoxia; however, this effect is lost at 7 days and instead 300μM levodopa inhibits ΔΨ_M_. Photomicrographs show TMRM staining in DRG soma treated with 0μM levodopa (left) or 300μM levodopa (right) in hypoxia. Scalebar = 10μm, for both images. B) left: Beta III tubulin immunocytochemistry shows no effect of high-dose (300μM) levodopa on soma size; however, mean percent fluorescence for ATP5b (B, right), a mitochondrial marker, was reduced. C) No impact of levodopa is observed in the context of parkinsonism (rotenone) at 24hrs. D) At 7 days, the deleterious effect of 300μM levodopa is maintained in mild ΔΨ_M_ inhibition (1nM rotenone). Further, both 30μM and 300μM levodopa reduce ΔΨ_M_ caused by 10nM rotenone. No additive effects of levodopa are observed at stronger ΔΨ_M_ inhibition caused by 500nM rotenone. Data in A, B, C and D are shown as box plots of technical replicates with whiskers depicting 5-95% percentiles, black circles depicting remaining data points, lines depicting medians and “+” symbols depicting means for optimal interpretation[41–43]. Light and dark orange circles show experiment means. A, B right, C, D: dashed lines show 100% (mean of control cells not treated with levodopa or rotenone) for reference. *p<0.05, **p<0.01, ****p<0.0001.

There was no impact of levodopa in the context of parkinsonism (rotenone) at 24hrs (Figure 2 B, effect of levodopa F (2.5, 108.2) = 1.4, ns). However, high-dose levodopa maintained its detrimental effect on ΔΨ_M_ at 7 days in all rotenone conditions – indeed, 300μM reduced ΔΨ_M_ to the same degree as 500nM rotenone (Figure 2 C). Moreover, 30μM worsened ΔΨ_M_ in cells treated with 10nM rotenone (Figure 2C, effect of levodopa F (2.8, 561.2) = 30.0, p<0.0001; levodopa × rotenone interaction F (7.5, 514.2) = 7.9, p<0.0001). This suggests that levodopa, at concentrations observed in patients, can exacerbate reductions in ΔΨ_M_. Moreover, high-dose levodopa, alone, impairs mitochondrial membrane potential in sensory neurons.

The loss in ΔΨ_M_ may be due to impaired electron transport impeding ATP production, as suggested by previous data at high-dose levodopa[32]; however, mitochondrial load itself may be reduced due to impaired transport[51]. We investigated this in a separate series of experiments where we analysed ATP5b content in DRG soma incubated for 7 days in hypoxia and treated with 300μM levodopa or control. Although there was no change in soma size, based upon beta III tubulin, indeed, mean fluorescence for ATP5b was reduced, suggesting some loss of mitochondrial from DRG soma (area: Figure 2B left unpaired ttest, t=1.71, df=254, ns; ATP5b fluorescence intensity Figure 2B right ttest, t=6.4 df=254, p<0.0001).

#### 3.2.2 Chronic levodopa initially increases and then ameliorates oxidative stress, at concentrations observed in vivo

Loss of mitochondrial membrane potential can lead to an increase in oxidative stress due to the failure of electron transport across complex I, in the case of rotenone, and subsequent transfer of the electron to acceptor molecules. We used dihydroethidium (DHE), which fluoresces when it reacts with reactive oxygen species.

After 24 hours treatment with levodopa only, no oxidative stress was observed, in keeping with the lack of change in ΔΨ_M_ at this timepoint (supplementary file S1, Figure S1 A; no effect of levodopa F (2.2, 93.0) = 2.9, ns; levodopa × condition F (3, 127) = 1.4, ns). In parkinsonian (rotenone-treated) sensory neurons cultured in normoxia, the highest dose of levodopa tended to reduce oxidative stress at 300μM (supplementary file S1 Figure S1 B: effect of levodopa F (2.5, 191.1) = 8.8, p<0.0001, levodopa × rotenone interaction F (9, 228) = 4.1, p<0.0001), in contrast to previous data[32]. In hypoxia, levodopa had no robust effects (effect of levodopa F (2.3, 176.3) = 4.6 p<0.01, levodopa × rotenone interaction F (9, 227) = 2.2 p<0.03). Thus, levodopa induced mild inconsistent effects on oxidative stress at 24hrs in parkinsonian sensory neurons.

After 7 days of treatment, levodopa alone caused a mild increase in oxidative stress, correlating with the loss in ΔΨ_M_ at this levodopa concentration and timepoint (Figure 3 A; Brown-Forsythe ANOVA test, F (3.0, 371.7) = 9.1, p<0.0001). The lowest level of rotenone (1nM) did not change ΔΨ_M_ and, in this context, levodopa also increased oxidative stress. However, at moderate levels of rotenone-induced inhibition of ΔΨ_M_ (10nM, 100nM), levodopa tended to reduce oxidative stress – this is not unexpected as the catechol group in levodopa may accept electrons[32] (Figure 3 B: levodopa × rotenone interaction F (9, 1027) = 12.26, p<0.0001). However, this ability may be overwhelmed at high levels of mitochondrial rotenone-mediated inhibition (Figure 3 B: 500nM).

**Fig 3.**
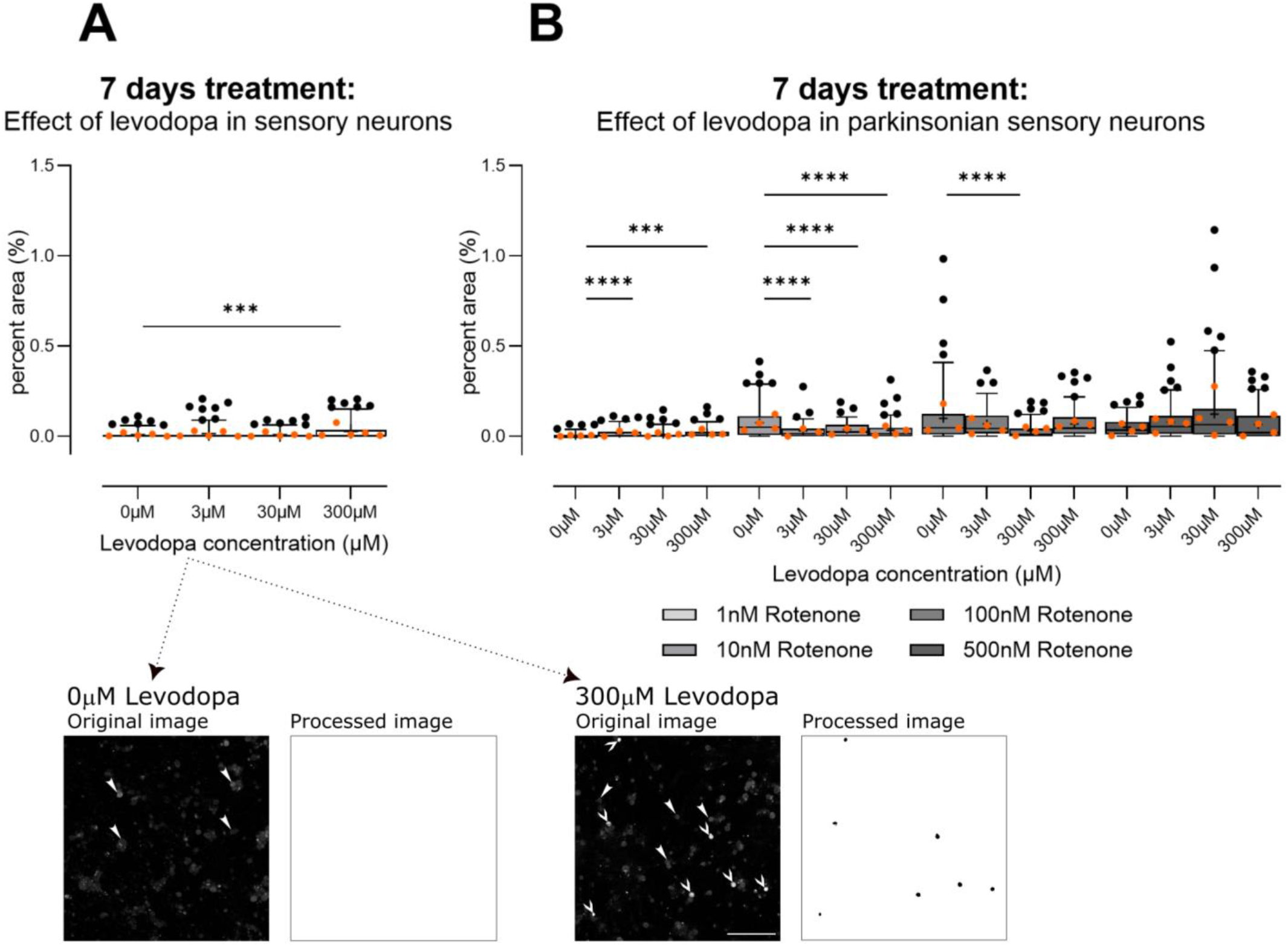
Effect of levodopa on oxidative stress in primary sensory neurons (DRGs) cultured in hypoxia. A) Alone, only the highest concentration of levodopa (300μM) induces mild oxidative stress. B) In the context of parkinsonism (rotenone), levodopa increases oxidative stress at 1nM rotenone. At moderate mitochondrial inhibition (10nM, 100nM), levodopa tended to reduce oxidative stress, possibly due to its ability to accept electrons[32] and this effect is lost at 500nM rotenone. Data are technical replicates and are shown as box plots with whiskers depicting 5-95% percentiles, black circles depicting remaining data points, lines depicting medians and “+” symbols depicting means for optimal interpretation[41–43]. Light and dark orange circles show means for each experiment. ***p<0.001, ****p<0.0001 post-hoc tests, following ANOVA, as discussed in text. Photomicrographs show example original images of cells treated with 0μM levodopa for 7 days with corresponding processed image or cells treated with 300μM levodopa for 7 days, and its corresponding process image. Arrowheads in photomicrographs point to DRG soma negative for oxidative stress (e.g., 0μM levodopa). Arrows in photomicrographs point to DRG soma positive for oxidative stress. Photomicrographs are not modified. Image processing picked out only DRG soma that contained reactive oxygen species. Scalebar = 200μm, for all images.

#### 3.2.3 Levodopa stabilizes tubulin at concentrations observed in vivo

Interestingly, levodopa has been suggested to incorporate into tubulin as a false amino acid and stabilize it, thereby preventing organelle transport[51] or interfering with protein degradation[52]. We therefore examined immunoreactivity for beta III tubulin, which is only found in neurons. Following 24hrs, both 3μM and 30μM levodopa led to an increase in percent area immunoreactive for beta III tubulin, with 300μM having no effect (Figure 4A, Brown-Forsythe ANOVA test F (3.0, 134.9) = 12, p<0.0001). In the context of parkinsonism, i.e., rotenone treatment, levodopa similarly led to increased immunoreactivity, particularly at 30μM, although 300μM increased immunoreactivity at 1nM and 10nM rotenone (Figure 4B; effect of levodopa F (2.4, 266.3) = 15.8, p<0.0001; levodopa × rotenone interaction F (9, 333) = 2.3, p<0.02). We then examined fluorescence at the level of individual neurites. Again, 30μM levodopa led to an increase in fluorescence in the context of levodopa alone (Figure 4C Brown-Forsythe ANOVA test, F (3.0, 135.7) = 6.6, p<0.001). In the context of parkinsonism, there was a trend towards dose-dependent increase in fluorescence, with 30μM and 300μM increasing fluorescence significantly (Figure 4D, effect of levodopa, F (2.5, 281.3) = 13.5, p<0.0001, levodopa × rotenone interaction F (9, 333) = 2.4, p<0.02).

**Fig 4.**
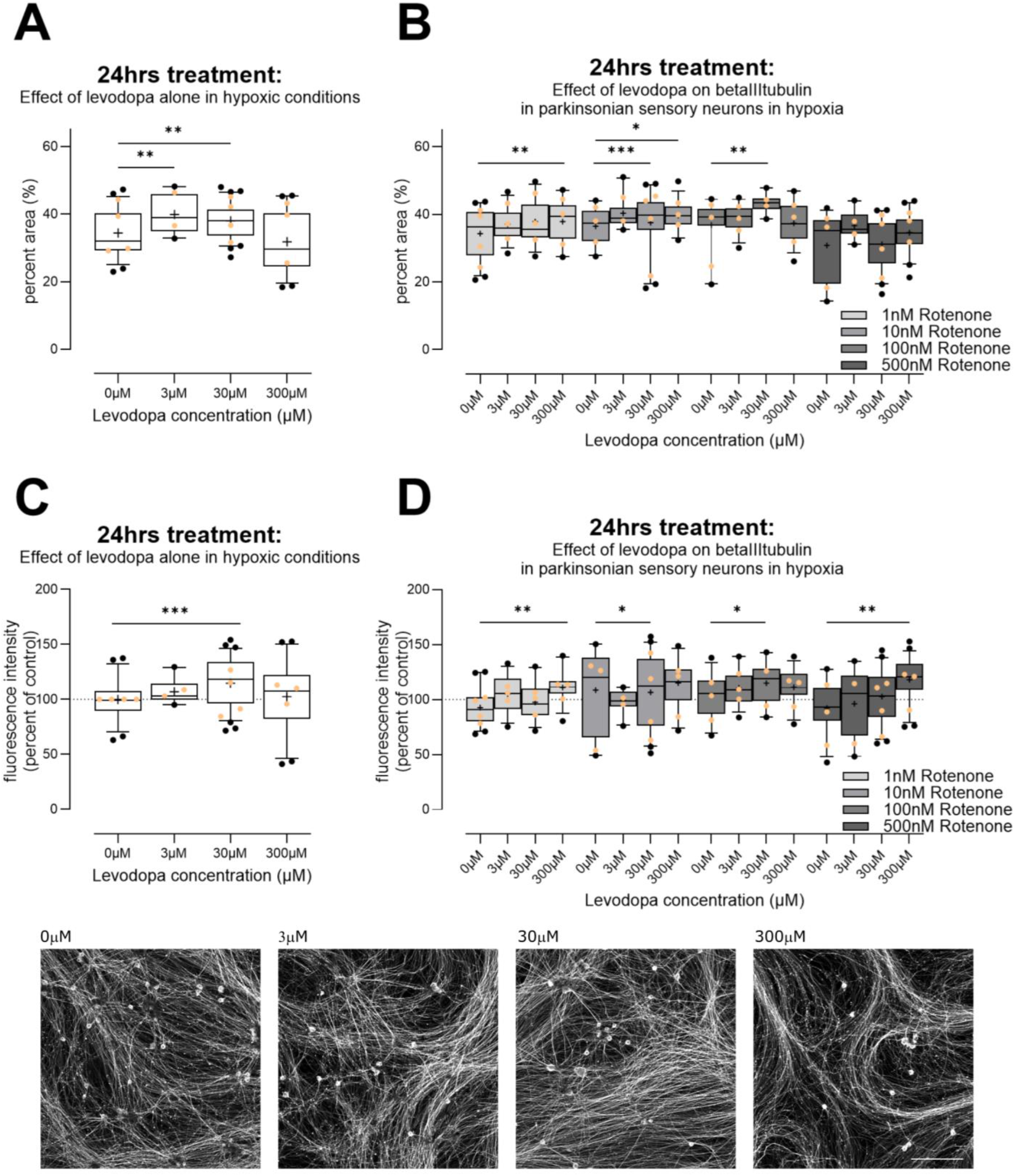
Effect of levodopa on beta III tubulin in primary sensory neurons (DRGs) cultured in hypoxia. A) Alone, 3μM and 30μM levodopa increased percent area immunoreactive for beta III tubulin. B) In the context of parkinsonian sensory neurons (treated with rotenone), levodopa tended to increase percent area positive for beta III tubulin at 30μM and 300μM. C) We examined fluorescence intensity for beta III tubulin, at the level of individual neurites in cells. Cells treated with levodopa only again showed increased fluorescence at 30μM. D) In the context of parkinsonian sensory neurons (treated with rotenone), 30μM and 300μM levodopa tended to increase fluorescence for beta III tubulin. Data are technical replicates shown as box plots with whiskers depicting 5-95% percentiles, black circles depicting remaining data points, lines depicting medians and “+” symbols depicting means for optimal interpretation[41–43]. Light orange circles depict experimental means. *p<0.05, **p<0.01, ***p<0.001 post-hoc tests, following ANOVA, as discussed in text. Photomicrographs show example original images of cells treated with 0μM, 3μM, 30μM, 300μM levodopa, only. Photomicrographs are not modified. Scalebar bottom right = 200μm, for all images.

#### 3.2.4 Levodopa reduces lysosome content at concentrations observed in vivo

Having established that 30μM levodopa exacerbates ΔΨ_M_ loss and may stabilize beta III tubulin in the context of parkinsonism, we then examined lysosomes because mitochondrial impairment plays a strong role in inducing lysosome biogenesis[53] and microtubules are critical for lysosome motility [54] but also because impairment of lysosome function is strongly implicated in PD[55]. For these experiments, we examined effects of levodopa alone, in hypoxic conditions.

In primary DRGs, despite the reduced mitochondrial membrane potential observed in DRGs at high-dose levodopa (Figure 2A), we observed reduced lysosomal content (Figure 5A, unpaired ttest, t=3.8, df=200, p<0.001), and a reduction in the proportion of lysosomes that were acidic (Figure 5B, unpaired ttest t=3.5, df=200, p<0.001).

**Fig 5.**
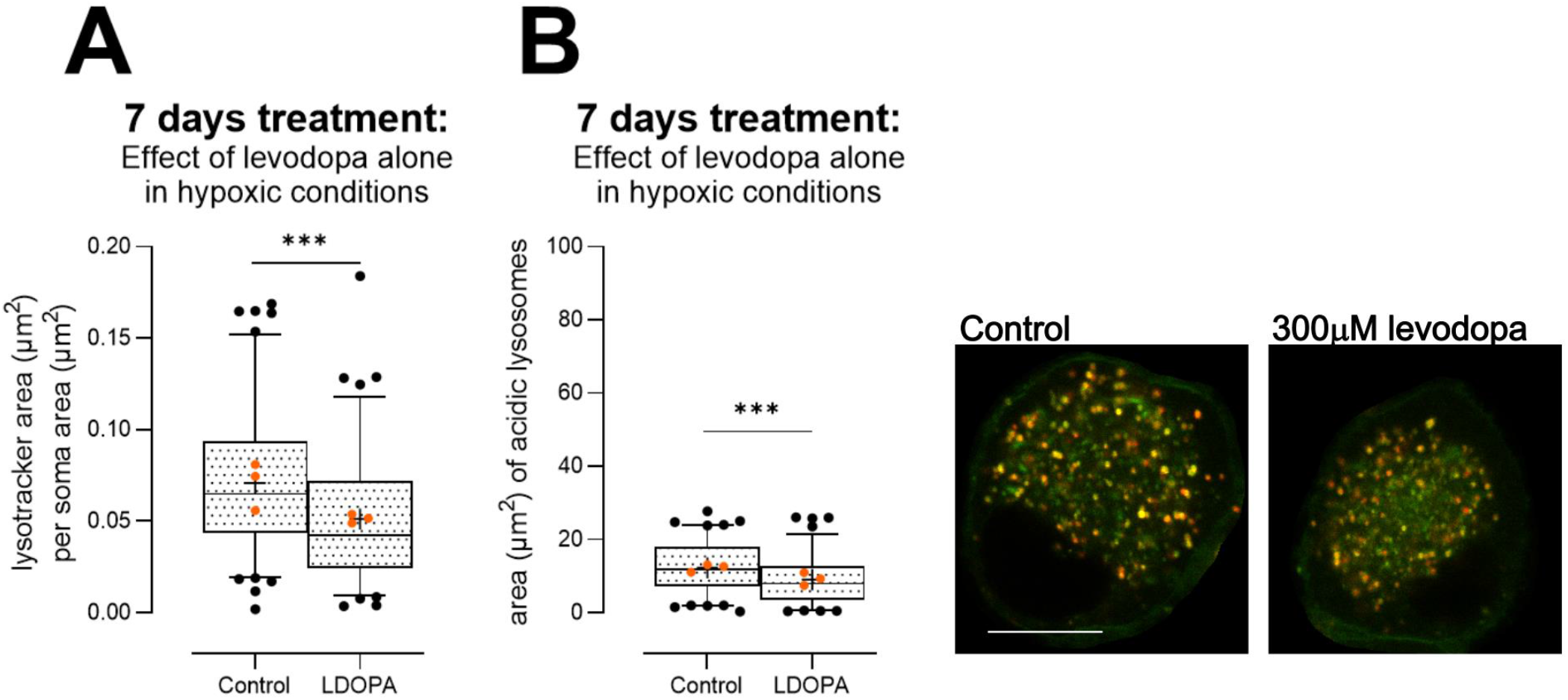
Effect of levodopa alone on lysosome content and acidity in primary DRGs cultured in hypoxia for 7 days. A) 300μM levodopa reduced lysosome content in DRG soma. B) Acidity of lysosomes was impaired by 300μM levodopa, because less lysosomes labelled with Lysotracker were co-labelled with Lysosensor. Data are shown as box plots with whiskers depicting 5-95% percentiles, black circles depicting remaining data points, lines depicting medians and “+” symbols depicting means for optimal interpretation[41– 43]. ****p<0.0001. Orange circles show experiment means. Photomicrographs depict control (left) and levodopa-treated (right) soma. Scalebar = 10μm, for both images.

We investigated these findings further in the 50B11 immortalized sensory neuronal cell line[56]. We examined in combination entacapone (1μM, [37]), which is used as an inhibitor of catechol-o-methyl-transferase in patients to prevent metabolism of levodopa to 3-o-methyldopa and subsequent formation of homocysteine. This enzyme is expressed by 50B11 cells[57]. We also examined homocysteine itself (20μM), used at concentrations that are observed in patients (10-20μM[38,39]).

We first examined sizes of all lysosomes. As this resulted in very large group sizes, we extracted 20000 lysosomes at random from the full dataset for each condition, to ensure an equal number for comparisons for the ANOVA. Both levodopa and entacapone affected lysosome size overall (effect of levodopa 2-way ANOVA, F (2, 119994) = 119.43, p=1.5e-52; effect of entacapone F (1, 119994) = 29.38, p=5.9e-8) and there was a strong interaction (F(2, 119994) = 16.1, p=1e-7). We then analysed the complete datasets using non-parametric 1-way analyses. 30μM levodopa, alone, caused a mild 3% increase in median size of lysosomes. However, 300μM levodopa had a strong effect and median size was reduced by 11% (Table 1, Kruskal Wallis anova H (3) = 175, p<0.0001). Treatment with entacapone exacerbated these effects with 30μM levodopa now causing a 6% loss in size and 300μM resulting in a 14% loss in median size compared with cells treated with entacapone only (Table 1, Kruskal Wallis anova H (3) = 419.3, p<0.0001). Homocysteine increased median lysosome size (homocysteine versus control-treated cells; Table 1, Mann Whitney U = 449847789, p<0.0001). Entacapone also showed no effect (control versus control + entacapone Mann Whitney U = 437670689, ns). Lysosome sizes per experiment (median ± 95% CIs) are shown in Table S1. Thus, lysosome content is reduced by levodopa at concentrations observed in the patient.

**Table 1.** Population estimates of median individual lysosome size^1^ in 50B11 cells treated for 24hrs in hypoxia.

| Condition | Median (95% CIs<sup>2</sup>) | Dunn's multiple comparisons test (versus Control) | Dunn's multiple comparisons test (versus Control + Entacapone) | Mann Whitney U test compared with control cells (no entacapone) |
| --- | --- | --- | --- | --- |
| Control | 35 (34-35) | - |  |  |
| 30µM levodopa | 36 (35-36) | * |  |  |
| 300µM levodopa | 31 (30-32) | **** |  |  |
| Control + 1µM entacapone | 35 (35-36) |  | - |  |
| 30µM levodopa + 1µM entacapone | 33 (32-33) |  | **** |  |
| 300µM levodopa + 1µM entacapone | 30 (29-30) |  | **** |  |
| Homocysteine 20µM | 38 (37-38) |  |  | ns |

<sup>1</sup>In pixels, as is standard for CellProfiler output<sup>2</sup> 95% confidence intervals of median
\*\*\*\*p<0.0001
ns non-significant

As Lysotracker is sequestered in acidic environs, we also examined mean fluorescence of lysosomes. We observed very mild effects. At 30μM levodopa, median fluorescence was increased by 2% from control-treated cells and at 300μM levodopa, median fluorescence was reduced by 5% (Table S2; Kruskal Wallis anova H (3) = 1879, p<0.0001). In the context of entacapone, median fluorescence was not changed by 30μM levodopa but was reduced by 3% by 300μM levodopa (Table S2; Kruskal Wallis anova H (3) = 614.0, p<0.0001). Homocysteine caused a mild increase in fluorescence (compared with control-treated cells, Mann Whitney U = 456646454, p<0.0001), as did Entacapone (compared with control-treated cells, Mann Whitney U = 422366710, p<0.0001). Thus, together, our data in 50B11 cells may reflect an absolute loss of lysosomes or a loss of acidic lysosomes. However, we note that our data in primary DRG soma (Figure 5) showed both a loss in lysosomes and remaining lysosomes were less acidic, thus it is likely that the same is occurring in 50B11 cells. Importantly, we show a loss of lysosomes at concentrations of levodopa, and entacapone, observed in the patient.

These analyses are on the basis of individual lysosome size, and had a very large N, which may have contributed to the very robust statistical differences that we observed. We then moved to a separate and more conservative per-cell based analysis by examining lysosome content at increasing distances from nuclei. As shown in Figure 6, we indeed observed a large reduction in total lysosome content at 300μM levodopa irrespective of entacapone (Figure 6A, effect of levodopa F (2, 146) = 5.8, p<0.01, distance × treatment F (8, 584) = 5.2, p<0.0001; Figure 6B, effect of levodopa, F (2, 179) = 13.08, p<0.0001, distance × treatment F (8, 716) = 11.8, p<0.0001). Critically, the addition of entacapone to protect levodopa resulted in a reduction in lysosome content caused by 30μM levodopa (Figure 6B). We also analysed these data using linear regression, and indeed, in the context of entacapone, the slope of increase in lysosome content within increasing distance from nucleus was reduced by both 30μM levodopa (F (1, 621) = 6.1, p<0.02) and 300μM levodopa (F (1, 636) = 15.4, p<0.0001).

**Fig 6.**
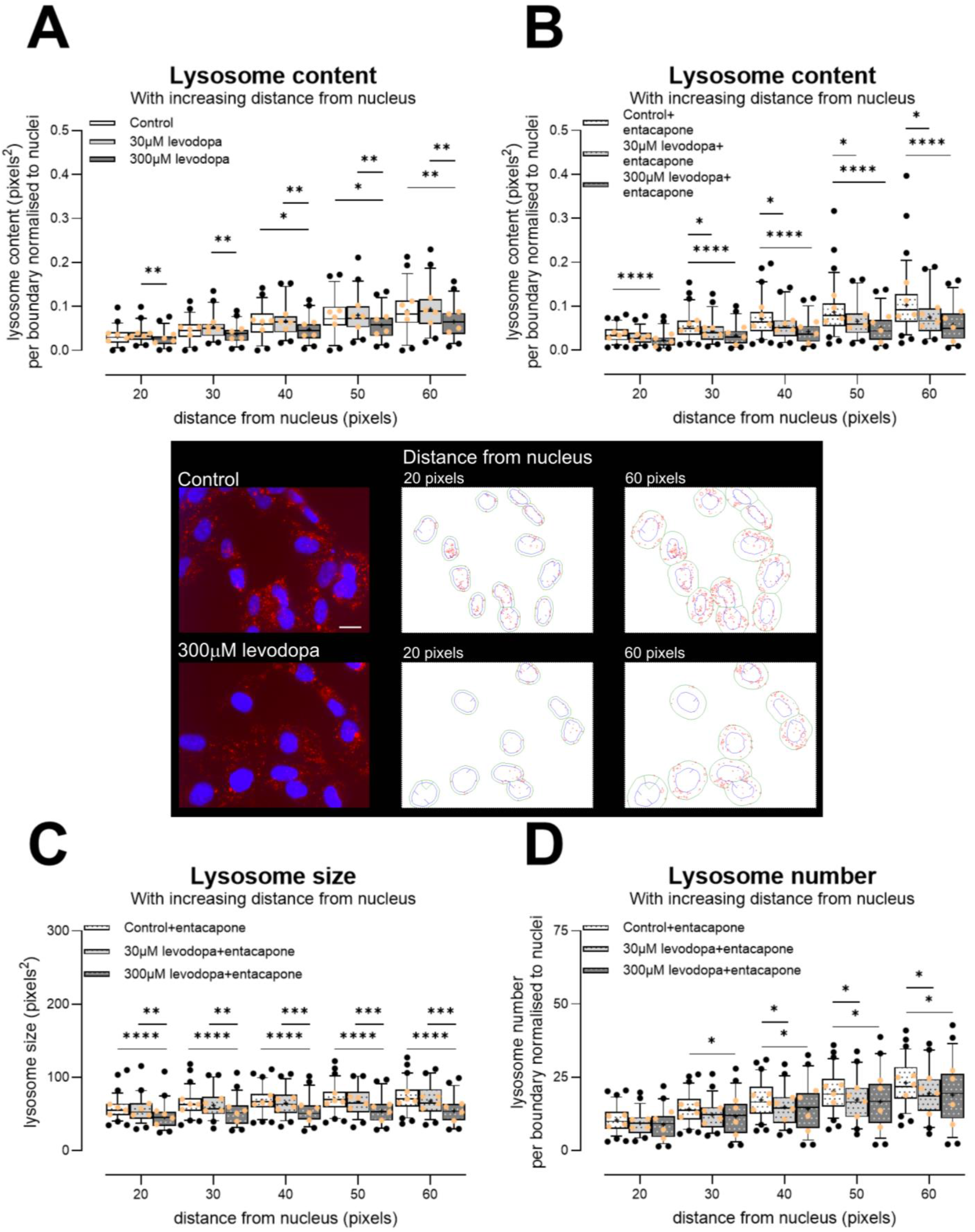
Effect of levodopa on lysosomes in the 50B11 immortalised sensory cell line, treated and incubated in hypoxia for 24 hrs. A) 300μM levodopa reduced lysosome content in comparison to control-treated cells and cells treated with 30μM levodopa. B) Both 30μM and 300μM levodopa reduced lysosome content when cells are treated with entacapone. Photomicrographs show example images of cells treated with 0μM or 300μM levodopa for 24 hrs in hypoxia. Photomicrographs are not modified except for brightness and contrast. Scalebar = 20μm, for both photomicrographs. Outlines show examples of CellProfiler segmentations of lysosomes (red) at 20 pixels and 60 pixels from individual nuclei (blue). Note: For segmentation, the entire nucleus was required to be within the field of view, and not to touch the image border. The green lines show the appropriate boundaries. C) Individual lysosome size per boundary, showing that in the context of entacapone, 300μM levodopa in particular reduces lysosome size. D) Number of lysosomes per boundary. In the context of entacapone, both 30μM and 300μM levodopa reduce lysosome number. Data in graphs are shown as box plots with whiskers depicting 5-95% percentiles, black circles depicting remaining data points, lines depicting medians and “+” symbols depicting means for optimal interpretation[41–43]. *p<0.05, **p<0.01, ***p<0.001, ****p<0.0001. Light orange circles show experiment means. Entacapone = 1μM [37].

With these smaller datasets, individual lysosome size was reduced in particular by 300μM levodopa, in the context of entacapone (Figure 6C; F (2, 179) = 15.0, p<0.0001, distance × treatment F (8, 716) = 2.7, p<0.01). Moreover, the number of lysosomes was reduced by both 30μM and 300μM levodopa, in the context of entacapone (Figure 6D; F (2, 179) = 4.5, p<0.05, distance × treatment F (8, 716) = 5.4, p<0.0001). When we compared control-treated cells to homocysteine-treated cells, there was slightly greater content (Figure S2A) and lysosome number (Figure S2C). In these smaller datasets, there was no change in size caused by homocysteine (Figure S2B). These data again suggest that metabolism to homo-cysteine does not underlie the deleterious effects of levodopa. Entacapone itself had little effect (control versus control + entacapone lysosome content effect of treatment F (1, 126) = 2.3, ns; control versus control + entacapone lysosome size effect of treatment F (1, 125) = 1.2, ns). Thus, our data show in two separate cell types that even when using conservative analyses, concentrations of levodopa observed in the patient reduce lysosome content and number.

## 4. Discussion

Levodopa is undoubtedly an important drug – it is on the WHO list of essential medicines[58] and is preferred among patients and neurologists for the treatment of the bradykinesia of PD [7–10]. However, at high doses it has been shown to exacerbate or initiate symptoms of peripheral neuropathy in patients [10,22]. It has been shown to impair nerve conduction in as little as one month in humans with effects worsening at later timepoints [31]. Additionally, oral levodopa dose correlates with loss of intraepidermal nerve fibres (IENFs) in skin [29].

With regard to the mechanism underlying these effects, levodopa is metabolized by the enzymes dopa decarboxylase and catechol-o-methyl transferase (COMT) [59]. The conversion by COMT leads to the formation of 3-o-methyldopa and homocysteine, both of which form reactive oxygen species and are toxic in vivo and in vitro [60–62]. Increased levels of homocysteine have repeatedly been found in PD [63–66] and are associated with cognitive decline in PD [64,66]; however, whether this plays a role in sensory deficits is unknown. Vitamin B12 is a co-factor in the catabolism of homocysteine and recommendations have been made that PD patients are treated with B vitamins including B12 and B6 to prevent peripheral neuropathy[10]. However, other mechanisms have also been postulated: high-dose levodopa has been linked to reduced mitochondrial respiration in immortalised mesencephalic cells and cancer cell lines [32,67], and levodopa is incorporated as a false amino acid into proteins, which inhibits proteolysis [52]. Incorporation of the false amino acid into tubulin may interfere with organelle transport[51].

Here, we indeed noted depolarization of ΔΨ_M_ at high chronic concentrations (300μM) in primary sensory neurons, which is in keeping with these previous data on high-dose levodopa in cancer cell lines [32]. Extending previous data that examined short timepoints[32], the impairment in ΔΨ_M_ caused by high-dose levodopa alone was associated with increased oxidative stress.

We further showed that levodopa causes hyperpolarization of the mitochondrial membrane potential (ΔΨ_M_) at low doses (30μM). However, in parkinsonian sensory neurons treated chronically with rotenone, low-dose levodopa exacerbated loss of ΔΨ_M_, suggesting that even low-dose levodopa can impair mitochondrial function in conditions of pre-existing mitochondrial dysfunction. We note that our low-dose levodopa mimics concentrations observed in the patient [45–50]. In the context of parkinsonism, low-dose levodopa reduced oxidative stress, presumably due to the limited ability of the levodopa molecule to scavenge free radicals[32]. Again at concentrations observed in the patient, levodopa stabilized beta III tubulin content over the short term, an important finding as previous data suggested that very high concentrations were required for this outcome[51].

Finally, and critically, despite the impairment of mitochondria at high doses, which would be expected to enhance lysosome biogenesis[53], we show that high-dose levodopa leads to *reduced* lysosomal content, and reduced acidic lysosomes in primary sensory neurons. These results were reproduced in 50B11 cells, which are immortalized sensory neurons [56], where individual lysosome size was reduced per cell at the highest concentration of levodopa. Moreover, when we co-treated our cells with low-dose levodopa and entacapone to protect levodopa from metabolism, effects on lysosomes were additive in 50B11 cells, with low-dose levodopa reducing content and number of lysosomes. Interestingly, homocysteine tended to have the opposite effect, causing a weak increase in content and number. These data suggest again that levodopa underlies the deleterious effects rather than homocysteine. Entacapone is an inhibitor of the enzyme COMT, which metabolises levodopa and leads to the formation of homocysteine, and that COMT is expressed by 50B11 cells[57]. Entacapone had very little effect on lysosomes.

Lysosome number and size are constantly changing within the cell as a homeostatic response to, e.g., nutrient availability. During nutrient deprivation and induction of autophagy, lysosome size typically increases and number decreases[68], and they also tend to sequester to the perinuclear area[69]. In lysosomal storage diseases, the inability to degrade substrates causes a massive increase in lysosomes size, and lysosomes are also increased in size in response to TMEM106B overexpression[68]. However, loss of the V-ATPase decreases lysosome number and size[68,69]. Interestingly, this proton pump is critical for lysosome acidification, and that is dependent upon energy levels of its activity[70].

Given the strong impact of translationally relevant concentrations of levodopa on tubulin that we observed here and that previously, very high levels of levodopa have stabilized tubulin and impaired mitochondrial transport[51,71], it is possible that lysosome content was impacted by a reduction in motility. However, this is unlikely to underlie the loss in acidity that we observed. Our data clearly show that already at concentrations observed in the patient, levodopa is deleterious towards lysosomes. We note that this low concentration did reduce oxidative stress in parkinsonian sensory neurons and stabilized tubulin over the short term; however, we do not maintain this as beneficial effects – the same concentration also impaired ΔΨ_M_ in parkinsonian sensory neurons.

Lysosomes are well established as being impaired in PD – the lysosome is the main method by which alpha synuclein is degraded[72]. Mutations in *GBA1* and *LRRK2* each increase the risk of PD and both GBA1 and *LRRK2* are important for lysosomal function as GBA1 encodes the lysosomal enzyme β-glucocerebrosidase[72] and LRRK2 may control lysosome levels within the cell[73]. Most PD patients are treated with levodopa and for very good reasons [7–10]. However, our data suggests that levodopa may contribute to peripheral neuronal dysfunction and may exacerbate lysosomal and mitochondrial impairment. Moreover, the lysosome releases vitamin B6 and B12 from their binding proteins[74,75], suggesting that levodopa may inhibit the desired effect of free B vitamins in patients.

## 5. Conclusions

Calls have been made to treat PD patients with B12[76] to alleviate the levodopa-mediated neuropathy. Here, in a highly translationally relevant series of experiments in sensory neurons, we showed that at concentrations observed in the patient, levodopa exacerbated mitochondrial impairment in parkinsonian cells, stabilized tubulin in parkinsonian cells and reduced lysosome content and acidity. Although B12 protects against homocysteine production, which is deleterious and associated with cognitive decline[77–79], homocysteine had very little effect on lysosome content or on lysosome acidity, outcomes that were affected by levodopa.

## Supplementary Materials

Supplementary file S1. Please also see Data availability statement below.

## Author Contributions

Conceptualization: OJO, MAH; methodology: MAH; formal analysis: OJO, AEA, AA, SH, MAH; resources: MAH; data curation: MAH; writing—original draft preparation: OJO, MAH; writing—review and editing: OJO, AEA, SH, AA, MAH; visualization: OJO, AEA, AA, SH, MAH; supervision: MAH; project administration: MAH. All authors have read and agreed to the published version of the manuscript.

## Institutional Review Board Statement

Not applicable.

## Data Availability Statement

Research data are available at https://doi.org/10.23673/re-477

## Acknowledgments

We thank Ulla Peterson for excellent technical assistance with primary cultures. 50B11 cells were a kind gift of Prof. Ahmet Höke of Johns Hopkins University, USA.

