## Supplementary file S1 for "Levodopa impairs lysosomal function in sensory neurons in vitro"

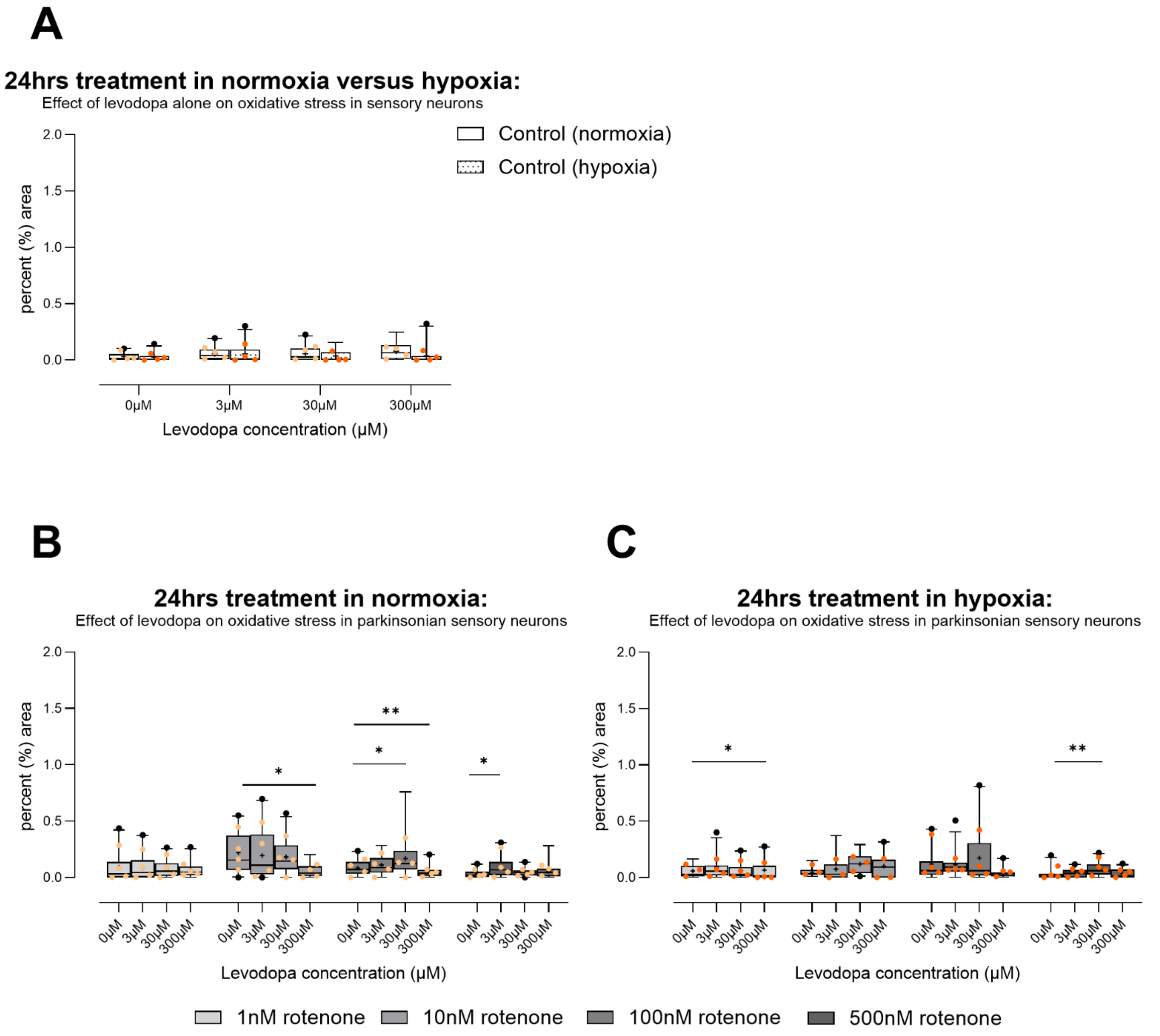


Figure S1 Effect of levodopa on oxidative stress in primary sensory neurons (DRGs) cultured in normoxia or hypoxia for 24 hours. A) Cells treated with levodopa alone for 24hrs show no oxidative stress whether in normoxia or hypoxia. B) In the context of parkinsonian (rotenone-treated) sensory neurons, 300µM levodopa tended to reduce oxidative stress in normoxia. C) Levodopa had no consistent effects in hypoxia. Data are technical replicates shown as box plots with whiskers depicting 5-95% percentiles, symbols depicting remaining data points, lines depicting medians and “+” symbols depicting means for optimal interpretation[1–3]. Light and dark orange symbols show experimental means (N=4 for normoxia and for hypoxia). *p<0.05, **p<0.01.

Table S1 95% confidence intervals of median lysosome sizes per experiment. Individual lysosome sizes (standardized, in pixels) in 50B11 cells treated for 24hrs in hypoxia.

| Biologic replicate | Control | Control + 1µM entacapone | 30µM levodopa | 30µM levodopa + 1µM entacapone | 300µM levodopa | 300µM levodopa + 1µM entacapone | Homocysteine 20µM |
|---|---|---|---|---|---|---|---|
| 1 | 33-36 | 34-37 | 30-33 | 27-29 | 24-26 | 21-23 | 35-38 |
| 2 | 33-36 | 32-34 | 35-38 | 33-35 | 33-35 | 30-31 | 36-37 |
| 3 | 35-37 | 38-40 | 36-38 | 36-38 | 31-33 | 32-33 | 41-44 |
| 4 | 32-34 | 31-34 | 34-36 | 29-31 | 28-31 | 27-29 | 35-36 |

Table S2 Median fluorescence of individual lysosomes in 50B11 cells treated for 24hrs in hypoxia*.

| Condition | Control<br><br>Median (95% CI of median) | With entacapone<br><br>Median (95% CI of median) |
|---|---|---|
| Control | 0.95 (0.95-0.96) | 0.95 (0.95-0.95) |
| 30µM LDOPA | 0.97 (0.97-0.98) | 0.95 (0.95-0.95) |
| 300µM LDOPA | 0.90 (0.90-0.91) | 0.92 (0.92-0.92) |
| Homocysteine | 0.96 (0.95-0.96) | - |

*Fluorescence is normalised to mean of control-treated cells.

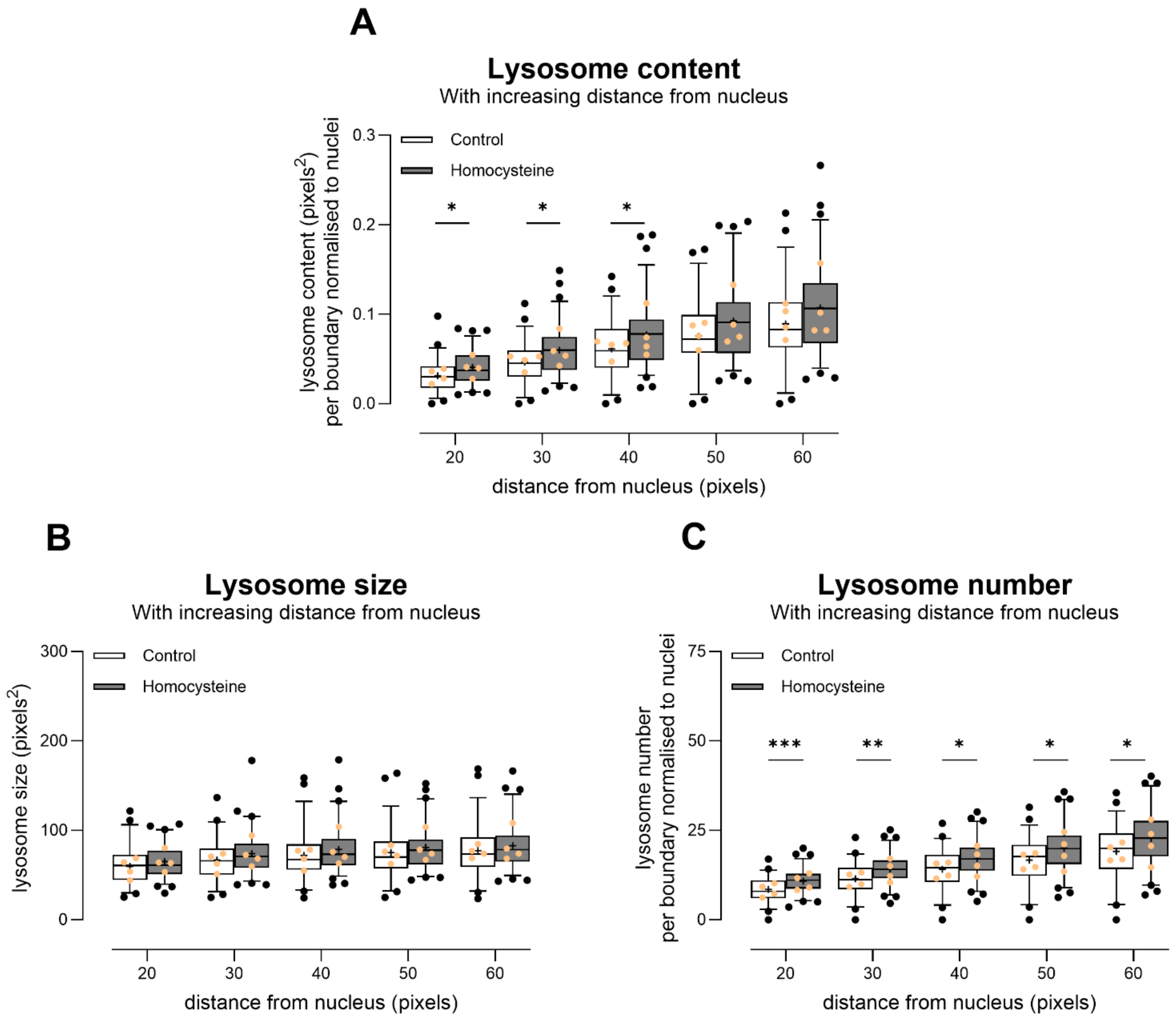


Figure S2 Lysosome content (A, effect of treatment F (1, 127) = 6.7, p<0.04) and number (C, effect of treatment F (1, 127) = 10.3, p<0.01) is increased slightly by 20µM homocysteine after 24hrs treatment in hypoxic conditions. There is no effect on individual lysosome size (B, effect of treatment F (1, 126) = 2.1, ns) in this small dataset. 50B11 cells (immortalised sensory neuronal cell line[4]) were habituated to hypoxic conditions and then treated and incubated for 24hrs in hypoxia. Data are shown as box plots with whiskers depicting 5-95% percentiles, black circles depicting remaining data points, lines depicting medians and "+" symbols depicting means for optimal interpretation[1–3]. Experiment means are superimposed as light-orange circles. * p<0.05, **p<0.01, ***p<0.001.
